# Wege: A New Metric for Ranking Locations for Biodiversity Conservation

**DOI:** 10.1101/2020.01.17.910299

**Authors:** Harith Farooq, Josue Anderson, Francesco Belluardo, Cristovao Nanvonamuquitxo, Dominic Bennett, Justin Moat, Amadeu Soares, Soren Faurby, Alexandre Antonelli

## Abstract

**Aim:** In order to implement effective conservation policies, it is crucial to know how biodiversity is distributed and one of the most widely used systems is the Key Biodiversity Areas (hereafter KBA) criteria, developed by the International Union for Conservation of Nature (IUCN). Here we develop a tool to rank Key Biodiversity Areas in a continuous scale to allow the ranking between KBAs and test this tool on a simulated dataset of 10 000 scenarios of species compositions of reptiles and mammals in eight locations in Mozambique.

**Location:** Mozambique, Africa

**Methods:** We compare the KBA criteria with four priorisation metrics (weighted endemism, extinction risk, evolutionary distinctiveness and EDGE score) to rank the biodiversity importance of eight sites with a randomly generated species composition of reptiles and mammals in Mozambique.

**Results:** We find that none of these metrics is able to provide a suitable ranking of the sites surveyed that would ultimately allow prioritization. We therefore develop and validate the “WEGE index” (Weighted Endemism including Global Endangerment index), which is an adaptation of the EDGE score (Evolutionarily Distinct and Globally Endangered) and allows the ranking of sites according to the KBA criteria but on a continuous scale.

**Main conclusions:** For our study system, the WEGE index scores areas that trigger KBA status higher and is able to rank their importance in terms of biodiversity by using the range and threat status of species present at the site. Prioritization may be crucial for policy making and real-life conservation, allowing the choice between otherwise equally qualified sites according to the KBA categories. WEGE is intended to support a transparent decision-making process in conservation.

## INTRODUCTION

In order to protect biodiversity and promote conservation, the decision-making process should be based more on scientific research and data, and less on expert judgement not supported by scientific studies (Sutherland et al., 2004). Threats to biodiversity such as conversion and degradation of natural habitats, and invasion by non-native species and overexploitation, have the potential of completely decimating biodiversity at local scales (Biofund, 2018; Mucova et al., 2018). Therefore, in recent years there has been an increased awareness of the value of protecting particular sites of high biological value, instead of focusing on large extensions of land (Butchart et al., 2012). Such decisions may ultimately determine whether biodiversity is preserved or lost. Thus, conservation planning should not only encompass the concepts of global conservation prioritization (Myers et al., 2000), but also include a more local-scale approach.

The Global Standards for the Identification of Key Biodiversity Areas (KBA) is an attempt to gather a consensus on the distribution of key biodiversity by highlighting sites that contribute significantly to the global persistence of biodiversity (IUCN 2004). The criteria and methodology for identifying KBAs was created by the IUCN World Commission on Protected Areas (IUCN, 2016). KBAs can vary considerably in size, and the criteria aim to address aspects of biodiversity operating from regional to relatively local scales. The categorization of areas is based on criteria such as presence and proportional inclusion of threatened species and ecosystems, species’ distribution ranges, ecological integrity and irreplaceability. However, indices that directly measure biodiversity such as species richness (SR), phylogenetic diversity (PD:Faith, 1992), weighted endemism (WE:Crisp et al., 2001) and phylogenetic endemism (PE:Rosauer et al., 2009) are not included in the KBA methodology.

Although most conservation prioritizations use a richness of already at-risk biodiversity locations (i.e., at-risk hotspots) in various forms. Few cases have the base information available on species richness to apply to practical site prioritization. Measures such as PD and PE introduce the evolutionary relations among species and minimize taxonomic uncertainty. All these indices contribute to the understanding of how and where biodiversity is distributed on a continuous scale, and should allow the ranking of individual sites under consideration for conservation. However, the accuracy of such indices is highly dependent on the quality and availability of data, making poorly sampled areas particularly hard to evaluate (Faith, 1992; Faith et al., 2004; Rosauer et al., 2009).

Although metrics may be useful in various ways in conservation, most of them fail to incorporate information on the threat status of the constituent species – the IUCN’s Red List Assessment parameter. One exception is the Evolutionarily Distinct and Globally Endangered (EDGE) score (Isaac et al., 2007), which combines one biodiversity index – Evolutionary Distinctiveness (ED) – with the threat category of species.

ED is a measurement of the branch lengths divided by the number of species within each clade. The EDGE score combines ED with values for species’ extinction risk in order to generate a list of species that are both evolutionarily distinct and globally endangered (‘EDGE species’). The EDGE score is however tailored to rank species rather than locations. Location scores may be computed as the sum of EDGE scores for all species at a site (Safi et al., 2013). However, this is not guaranteed to maximize conservation importance of individual sites, since the presence of widespread, critically endangered species produces higher EDGE scores than a vulnerable or endangered micro-endemic restricted to very few sites, which could rapidly go extinct if those sites are damaged. One example is the Atlantic bluefin tuna, which exists across a great part of the Atlantic Ocean, but nevertheless is considered an endangered species (Collette et al., 2011).

Without using biodiversity indices, systematic conservation planning is able to spatially prioritize areas for conservation attributing features to each area and by setting targets. One of the most common approaches is the use of the concept of irreplaceability (Pressey et al., 1994), so that irreplaceability scores are calculated for each conservation feature in each planning unit and range between 0 and 1 (Ferrier et al., 2000). Sites with values closer to 1 are considered irreplaceable if lost, while values closer to 0 are attributed to sites that in case of loss, targets may still be met by prioritizing other areas.

In this study we propose an index capable of ranking locations already meeting KBA criteria and compare its performance with WE, EDGE Score, ED and extinction risk (ER). Since the KBA methodology weights all species equally, irrespective of their evolutionary uniqueness, we excluded PD and PE from our analysis. We focus on two distinct vertebrate groups, reptiles and mammals, in which we generated 10 000 possible scenarios and tested the new index’s efficiency at prioritizing locations according to the KBA criteria. Our new spatially explicit index –WEGE (Weighted Endemism and Globally Endangered) is an adaptation of the EDGE score (Mooers et al., 2008), where we have replaced the phylogenetic component with an endemism score and is presented as a tool for attributing a continuous scale for the Global Standard of Identification of KBAs.

## METHODS

### Key Biodiversity Areas

Although the Global Standard for the Identification of Key Biodiversity Areas (KBA) (IUCN, 2016) has five main criteria and thresholds for the assessment, namely: A. Threatened biodiversity; B. Geographically restricted biodiversity; C. Ecological integrity; D. Biological processes; and E. Irreplaceability through quantitative analysis, only criteria A and B could be applied to our dataset consisting of a georeferenced species list. The full list of criteria and applicability in this study is provided in Appendix S1.

#### Study area and scenarios

In order to test the new index we propose here, we simulated communities of either reptiles of mammals in sets of eight location with hypothetical areas corresponding to 0.5 by 0.5 degrees (~2 500 km^2^) and with species numbers corresponding to the empirically observed from our field work (6, 7, 8, 8, 9, 9, 10 and 11 species). We simulated 10 000 combinations of species compositions for each location for both vertebrate groups. We restricted our analysis to species occurring in Mozambique and generated communities of random species known from within the country. We retrieved species occurrences from GBIF for all reptiles (https://doi.org/10.15468/dl.jwzffj) and mammals (https://doi.org/10.15468/dl.6hrjrx) and produced checklists for reptiles and mammals based on the species that had records in the country (reptiles - https://doi.org/10.15468/dl.fpyayo, mammals - https://doi.org/10.15468/dl.2wjwh9). We repeated the analysis for KBAs with 0.1 by 0.1 and 1 by 1degrees and the results can be found in the supplementary materials (Appendix S1) as well as the list of species used to simulate scenarios.

To calculate the distribution of species we rounded the GBIF records to 0.1 degrees, thus creating distribution maps composed of a sum of squares of ~100 km^2^.

To check whether a particular location would trigger KBA status, we restricted our analysis to three sub-criteria, A1a), A1b) and B1 of the KBA guidelines.

The criteria A1a) states that the site regularly holds ≥0.5% of the global population size AND ≥5 reproductive units of a CR or EN species;

The criteria A1b) states that site regularly holds ≥1% of the global population size AND ≥10 reproductive units of a VU species;

The criteria B1) states that Site regularly holds ≥10% of the global population size AND ≥10 reproductive units of a species;

We assumed the presence of ≥10 reproductive units whenever a species was present in a location.

To which we addressed by using the following conditions:

Presence of a CR or EN species with a distribution of 100 000 km^2^ or less (corresponding to a presence in one thousand 0.1-degree cells), presence of a VU species with a distribution of 10 000 km^2^ or less (corresponding to a presence in one hundred 0.1-degree cells) and presence of any species with a distribution of 1000 km^2^ or less (corresponding to a presence in 10 0.1-degree cells).

#### Biodiversity indices

To test whether we could use widely used biodiversity metrics to rank our locations, we calculated the scores of four indices: WE, EDGE score, ER and ED and compared the ranking of such metrics to our new index, WEGE.

Metrics such as EDGE, ER and ED, were calculated by summing the values of the species in each community randomly generated.

To compare the different ranking of the different metrics for each of the 10 000 scenarios we tested how often the different indices prioritize areas that trigger KBA status.

By using eight fictional locations, the number of areas triggering KBA status vary between 0 and 8 and the perfect ranking scores would vary between 1 for scenarios with 1 KBA and 36 for scenarios with 8 KBAs (1+2+3+4+5+6+7+8) (Appendix S1).

By comparing the distance between the obtained rankings from the different metrics and the perfect ranking score we are able to compare the performance of the different indices at ranking KBAs.

We compared the by calculating a ranking metric which we defined as (Obs-Min)/(Max-Min). Obs was the observed sum of the ranking of the sites scored as Kba (i.e. if a simulation had two KBAs and they are ranked as 2^nd^ and 4^th^ highest for the particular metric, Obs would be 2+4=6). Max and Min and are the highest and lowest possible rankings for the number of observed KBAs in a given simulation. The ranking score thus varied betwee 0 (perfect) and 1 (worst case scenario irrespective of the number of KbAs).

### The WEGE index

We sought a measure that would align its results with the IUCN’s KBA categorization of our locations. Since such measure has not yet been proposed to the best of our knowledge, we created an index capable of ranking locations in a continuous scale within the categories of the KBA.

The WEGE index proposed here is an adaptation of the EDGE score (Isaac et al., 2007) using the probability of extinction risk as in Mooers et al. (2008). The idea of the EDGE score is to measure biodiversity by taking into account both the evolutionary distinctness (ED) and the Probability of Extinction (ER) as an initial indication of “conservation priority” for species.

To calculate EDGE, the following formula is used in Mooers et al. (2008):

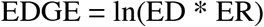

The WEGE index uses weighted endemism (WE) instead of evolutionary distinctness (ED) and just like EDGE, the probability of extinction (ER).

To calculate WEGE, we apply the formula:

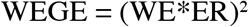

To calculate the WEGE index in any given site, we do a sum of the square of the partial weighted endemism value for each species multiplied by its probability of extinction value. In order to calculate the values for the WEGE index of all the locations in this study we created a package in R, available at devtools::install_github(‘harithmorgadinho/wege’).

We used the IUCN50 transformation for the ER as in (Davis et al., 2018), which scales the extinction risk over a 50-year period using the following extinction probabilities: LC = 0.0009, NT = 0.0071, VU = 0.0513, EN = 0.4276, CR = 0.9688.

The EDGE enables the ranking of species, rather than directly scoring areas, in regard to prioritization. The WEGE index, in contrast, allows the ranking of locations rather than individual taxa.

In order to give an extinction risk value to “DD” (Data Deficient) species, we assigned them with the same value as VU species following Bland et al. (2015) who concluded that 64% of mammals assigned to DD are at risk of extinction.

## RESULTS

By generating 10 000 species’ compositions of reptiles and mammals for eight fictional locations, we created eight different outcomes regarding the number of areas that could trigger KBA status Fig 1. A and Fig 2. A. For reptiles, due to the high number of range restricted species in the group, the simulation was able to create the eight possible scenarios, while mammals due to most species being widespread, there were less KBA trigger species to trigger KBA status, thus, no scenario with eight areas qualified as KBA was generated. In order to compare the ranking of KBAs between WEGE, WE, ER, ED and EDGE, we summed all the ranking scores, where the perfect ranking score takes the lowest possible value. Thus, the lower the sum of the ranking of the metrics, the shortest the distance to the perfect ranking. In both vertebrate groups, WEGE outperformed the other metrics, followed by WE, ER, EDGE and ED. In reptiles the difference between WEGE and WE was much smaller, 494.14 to 566.61 (Fig. 2 B) than in mammals, 89.09 to 435.36 (Fig. 2 B). This difference in performance is related to the fact that in mammals, unlike reptiles there are more widespread endangered species. These species have the potential of triggering KBA status and are weighted by WEGE, unlike WE, which doesn’t take into account the conservation parameter.

**Figure 1:**
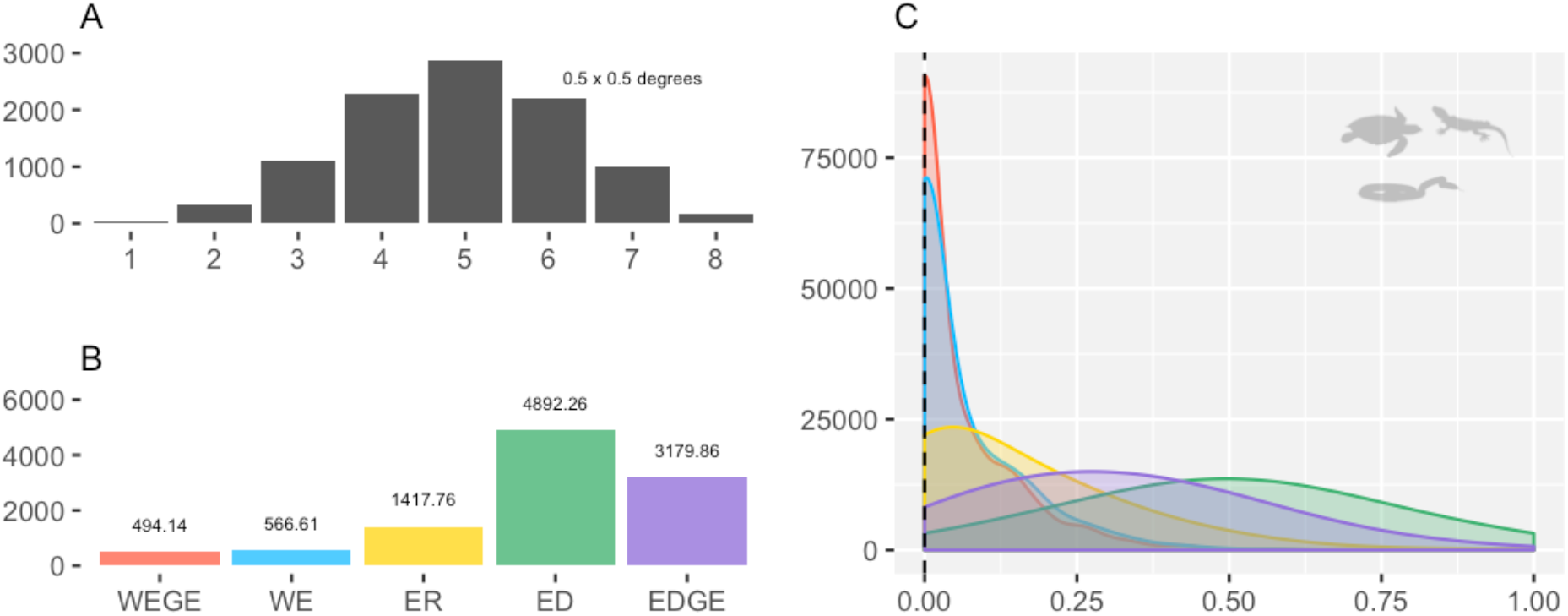
Number of areas triggering KBA status obtained by simulating 10 000 scenarios in reptile species’ composition. B. Indices combined sum for all scenarios. C. Frequency of scores normalized between different number of KBAs. The figure shows that WEGE outperforms the other indices by both getting a smaller overall sum (B) and by having a higher density of values closer to 0 (C).

**Figure 2:**
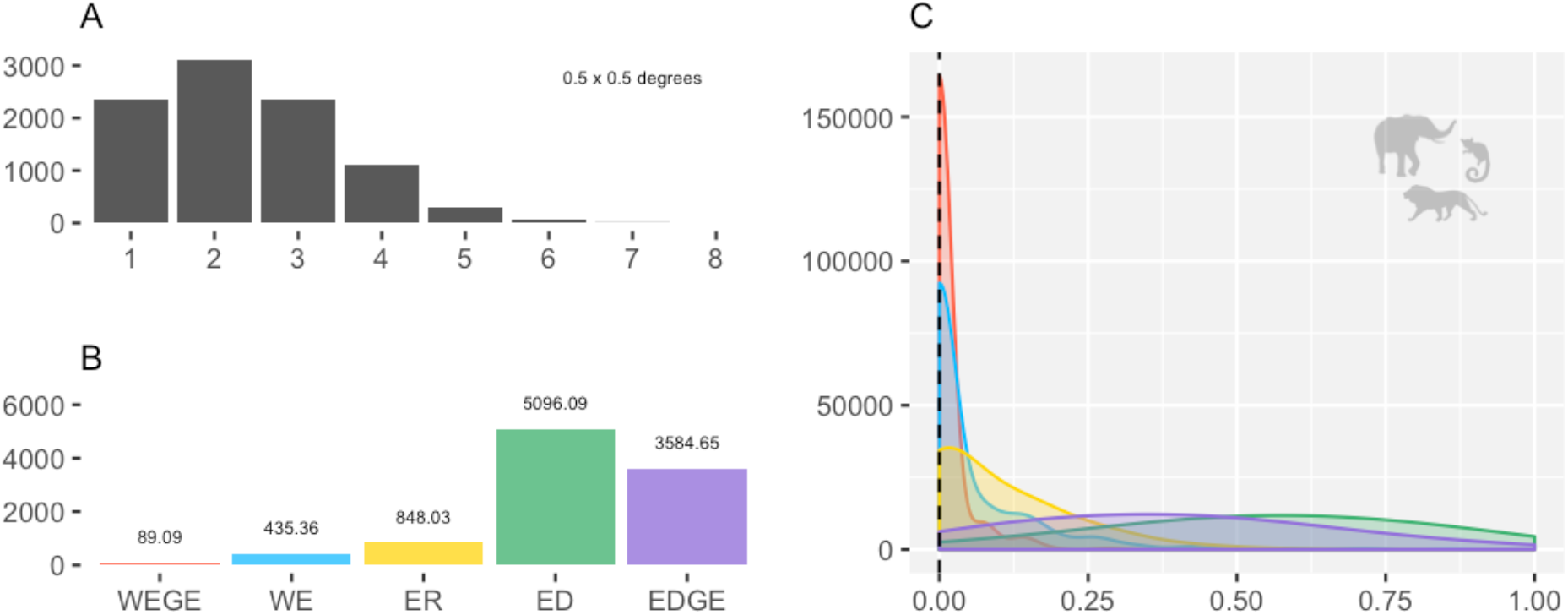
Number of areas triggering KBA status obtained by simulating 10 000 scenarios in mamal species’ composition. B. Indices combined sum for all scenarios. C. Frequency of scores normalized between different number of KBAs. The figure shows that WEGE outperforms the other indices by both getting a smaller overall sum (B) and by having a higher density of values closer to 0 (C).

In order to test the sparsity of the ranking scores we normalized the data where a score of 0 is the perfect score while the score of 1 is the worst. Both reptiles and mammals had most of their WEGE scores closer to 0 when compared to the other metrics (Fig 1. C and Fig 2. C). For both vertebrate groups tested in this study, reptiles and mammals, and for all KBAs sizes tested (0.1 by 0.1, 0.5 by 0.5 and 1 by 1 degrees), WEGE consistently outperforms WE, ED, ER and EDGE both in distance to perfect scores and in number of best rankings achieved. In addition, WEGE performed 6.4 times better at ranking KBAs than EDGE for reptiles and 40.2 times better for mammals. Thus, our study suggests that in order to rank KBAs in a continuous scale and using KBA’s criteria, WEGE performs substantially better than EDGE.

## DISCUSSION

### IUCN’s KBA and priorisation indices

The IUCN’s KBA uses a set of guidelines to check whether a particular site triggers a KBA status, unlike biodiversity metrics which attempt to quantify different spectra of biodiversity. Hence, different biodiversity metrics are expected to weight sites differently. The biodiversity of specific sites should arguably not be assessed by just summing the number of species existing in each location, but also taking into account other factors such as genetic diversity, distribution ranges or conservation status (Magurran, 1988; Barthlott et al., 1999). Otherwise, the presence of many widespread species producing a high SR would mask the importance of vulnerable or endangered micro-endemic taxa (restricted to very few sites).

The fact that SR and PD indices are known to be highly correlated with sampling effort (Bunge & Fitzpatrick, 1993; A. Rodrigues et al., 2005; A. S. Rodrigues et al., 2011; Tucker & Cadotte, 2013) advocates against their use in inconsistently and poorly sampled regions, compared to dense sampling which will in most cases show higher species diversity. In addition, SR and PD completely disregard the information on species range in their score, which is a strong predictor of extinction risks for species (Purvis et al., 2000) and one of the fundamental aspects of conservation prioritization and management of natural resources (Anderson, 1994; Myers et al., 2000; Roberts et al., 2002; Slatyer et al., 2007).

WE and PE are also expected to correlate with sampling effort, since new sets of records can only consolidate or increase the score but never decrease it (Lande, 1996; Nipperess, 2016), although this correlation seems to exist at lesser extent than in SR and PD (Soria-Auza & Kessler, 2008; Oliveira et al., 2016). But besides the sampling effort issue, the use of WE and PE in ranking areas might encounter additional problems. A benefit of PE is that for two recently diverged taxa, the vast amount of their evolutionary history is shared and it therefore matters very little if they are treated as separate species or not. This is critical for groups with large genera, which often comprise both widespread and range-restricted species as a result of species radiations. One example is the widely distributed skink of the genus *Cryptoblepharus*, which if analyzed through WE would score considerably higher compared with an analysis using PE. *Cryptoblepharus* is very widespread, with some species occurring from the eastern fringes of the Indo-Australian archipelago, Australia and Oceania, to the islands of the far Western Indian Ocean and adjacent parts of the African coast (Rocha et al., 2006). The WE index, in contrast, gives a weight of 1 for every species, which makes the index more vulnerable to taxonomic changes but guarantees the equal contribution of species within large genera.

In this study we tested the ranking of KBAs using WE, ER, ED, EDGE and our proposed index WEGE. Locations that have higher WE are areas in which the species composition contain species and more restricted ranges. Locations that score higher in ER are areas that contain more species with higher threat status. Locations with higher ED are areas that house species which have a higher evolutionary distinctiveness. Locations which score higher in terms of EDGE are areas that have a composition of species with both high evolutionary distinctiveness and threat levels. Finally, locations with higher WEGE scores, will be locations with a combinations of range restricted and threatened species.

Regarding our analysis, using 10 000 simulated scenarios of species compositions of reptiles and mammals, the WEGE index outperformed WE, ED, ER and EDGE both at overall sum of ranking scores and density of scores closer to the perfect score. The results were consistent for both vertebrate groups and KBA sites sizes tested. The second-best metric was WE, followed by ER, EDGE and in last place ED. Interestingly, our results show that using ER alone would be a more efficient way of ranking KBAs when compared to EDGE. Even though, both EDGE scores and the KBA initiative are focused on the preservation of biodiversity, according to our study they prioritize different sets of species.

The use of EDGE scores to rank sites is only expected to be efficient when the threats are plausibly mitigated by the protection of a site. Threatened species may be very widespread under two different scenarios, either they may live in very low population densities like in the case of tigers or they may be threatened by causes which are not geographic in nature and where protection of individual areas is of low importance such as in the case of of the Tasmanian devil. In both cases the species IUCN rank seems highly plausible but no single site will be as important for the protection of either of the two species as a site containing the majority of the range of a microendemic would be in the case of the Near threatened Mount Mabu Pygmy Chameleon.

### Suitability of the WEGE Index

The new index proposed here (WEGE) is capable of ranking locations in a continuous scale and matching the KBA status triggered by IUCN’s KBA. The WEGE index adds the component of conservation status of each species to the WE index. The internal logic of this metric is to combine conservation scoring of each species with a measure of the relative importance of the site in question for each species. This could also be achieved by combining a conservation score which incorporating evolutionary history such as e.g. PE rather than WE, but since KBA by design weigh all species equally irrespective of their evolutionary uniqueness we chose to select a measure with the same lack of taxonomic weighing. By incorporating WE in the EDGE score formula and creating the WEGE index, we obtained an index in line with the IUCN KBAs standards criteria compared to the WE, ED, ER and EDGE.

The WEGE index can be used either to find suitable candidates’ areas to be considered as KBA’s or as a mechanism of weighting the importance of biodiversity of particular KBA’s as well as areas outside KBA’s. Additionally, it uses a simpler methodology by employing only two metrics instead of a set of seven conditions (A1a - e and B1 and B2). Finally, WEGE can act as a complement in the process, by which, sites selected using IUCN’s KBA can now be ranked objectively according to their biodiversity importance.

Complementing the categorical ranking of locations can bring great advantages when prioritizing efforts with limited resources. IUCN’s criteria lack this aspect by attributing a binary system where one particular site either triggers KBA status or not. By using WEGE, we rank sites within the same category and enabling the decision-making process to be objective and transparent as possible. Getting conservation actions applied to any given area usually demands a great deal of effort, so sensitivity to removing sites from KBA status is less likely to be of high societal priority, but still, this methodology can highlight areas which even though they trigger KBA status, their score is low and might be on the cusp of losing their KBA status. KBA sites which are driven either by the presence of one threatened or range restricted species, will change if species become non-threatened or get their range considerably expanded. Consequently, lower performing WEGE sites have higher odds of losing their KBA status. One example that illustrates this scenario is the species *Cryptoblepharus ahli* Mertens, 1928, described by Mertens (1928), synonymized to the widespread species *Cryptoblepharus africanus* by Brygoo (1986) to later, based on a morphological examination of the species to be elevated to full species by (Horner & Adams, 2007). This species by itself meets the requirements for the Mozambican Island to trigger KBA status, regardless of its IUCN status, since it is at the moment an accepted species confined to a single small island. Further analysis of the genetics of this particular species will have an impact on the KBA status of this island.

### Limitations and challenges of the WEGE index

The two measures, EDGE and WEGE combine two clearly different and unrelated metrics, whilst WEGE makes use of species distribution and it’s IUCN status just as in the IUCN’s KBA. These two criteria are not independent since range size is one of the criteria for IUCN status. Importantly, however, the two criteria in WEGE clearly still measure distinct processes which for instance can be seen by the existence of widespread but endangered species like the already mentioned Bluefin tuna or highly restricted and least concern as the Mount Mabu Pygmy Chameleon. By combining the two we show that we get a better measure than solely relying on IUCN criteria or solely on WE.

Despite ranking locations according to the KBA’s guidelines, the WEGE index does not incorporate all the KBA’s criteria because it only uses georeferenced species lists to rank KBAs. Thus, information such as Ecological Integrity (criteria C), Biological Processes (Criteria D) and Irreplaceability Through Quantitative Analysis (Criteria E) may be subject to further attempts at complementing the WEGE index.

The last step of before proposing a particular site as a KBA requires an analysis of the manageability of the site in regards to its physical attributes such as forest cover limits or rivers and anthropogenic factors such as roads and existence of human settlements. The WEGE index in itself is not aimed at replacing this process, we believe this step to be of crucial importance and should be done case by case and involving local authorities. The aim of the WEGE index is to highlight and rank sites which should in the following step be scrutinized at a local level as in the KBA process or rank already existing KBAs.

### Final remarks

The idea of priorisation between KBAs foreseeing conservation policy has already been proposed (Pressey et al., 1994; Ferrier et al., 2000; Plumptre et al., 2019; Smith et al., 2019). Protection status, funding, irreplaceability by prioritization software (Plumptre et al., 2019) and systematic conservation planning (Smith et al., 2019) have been proposed to support the ranking of priority of areas. The results, although providing some kind of hierarchy between KBAs, still cluster KBAs in different categories, rather than scoring individual sites as allowed in WEGE. In systematic conservation planning, conservation practitioners must choose which conservation features should be used to represent biodiversity (Smith et al., 2019). WEGE represents a simple metric that encapsulates the biodiversity importance of a particular site, highlighting the same areas as the KBAs criteria while adding the component of continuous scale. Therefore, WEGE may also be used as a feature in systematic conservation planning.

The prioritization of areas in regard to biodiversity is complex. Different indices prioritize different areas. IUCN KBAs do not contemplate biodiversity indices in the decision-making process. However, for the case of reptiles and mammals in Mozambique, we found a correlation between the areas that would in theory trigger Key Biodiversity Area status and the WEGE Index.

Mozambique is a developing country that struggles to conciliate its rich biodiversity with the for the mining industry, and the high potential economic gain that could follow. The country also has one of the highest corruption levels in the world, and unbiased methods to quantify biodiversity are a crucial parameter for a transparent decision-making process in conservation. The selection of sites as KBAs is expected to have multiple uses, including conservation planning support and priority-setting at national and regional levels (IUCN, 2016). Therefore, the use of the WEGE index, allowing the ranking of key biodiversity areas is expected to by association support a transparent ranking of sites in regards to conservation.

## Supporting Information

Methods used for calculating indices, r packages used, KBA guidelines and raw data (Appendix S1), is available online.

## DATA AVAILABILITY STATEMENT

Data may be available from the authors upon request.

