## Supplementary Materials for "Wege: A New Metric for Ranking Locations for Biodiversity Conservation"

1. Appendix Sl

### Supplementary materials

Table S1: Reptile species list, Range in number of 0.1 degree units, ED: Evolutionary distinctiveness, ER: Extinction risk.

| status | species | Range | ED | ER |
|---|---|---|---|---|
| CR | Rhampholeon bruessoworum | 2 | 17.9105182 | 0.9688 |
| CR | Eretmochelys imbricata | 1615 | 41.4436829 | 0.9688 |
| DD | Scolecoseps boulengeri | 1 | 13.6560586 | 0.0513 |
| DD | Proscelotes aenea | 1 | 14.183756 | 0.0513 |
| EN | Atheris mabuensis | 2 | 8.94928558 | 0.4276 |
| EN | Rhampholeon gorongosae | 1 | 23.2422456 | 0.4276 |
| EN | Chelonia mydas | 2889 | 48.7286829 | 0.4276 |
| EN | Rhampholeon tilburyi | 3 | 17.6606998 | 0.4276 |
| EN | Cycloderma frenatum | 11 | 30.9951086 | 0.4276 |
| LC | Acanthocercus atricollis | 530 | 14.3176639 | 0.0009 |
| LC | Acontias aurantiacus | 18 | 11.4569221 | 0.0009 |
| LC | Acontias plumbeus | 154 | 12.5539558 | 0.0009 |
| LC | Afroablepharus wahlbergi | 226 | 10.8018431 | 0.0009 |
| LC | Afroedura langi | 12 | 13.824349 | 0.0009 |
| LC | Afroedura loveridgei | 4 | 19.4034067 | 0.0009 |
| LC | Afrotyphlops bibronii | 241 | 8.64549989 | 0.0009 |
| LC | Afrotyphlops fornasinii | 20 | 8.66027189 | 0.0009 |
| LC | Afrotyphlops mucruso | 83 | 8.39886601 | 0.0009 |
| LC | Afrotyphlops schlegelii | 134 | 8.51756579 | 0.0009 |
| LC | Agama aculeata | 590 | 11.2057552 | 0.0009 |
| LC | Agama armata | 143 | 11.2456838 | 0.0009 |
| LC | Agama hispida | 166 | 12.0943059 | 0.0009 |
| LC | Agama kirkii | 48 | 12.6284371 | 0.0009 |
| LC | Agama mossambica | 37 | 11.2715253 | 0.0009 |
| LC | Amblyodipsas concolor | 34 | 10.6589353 | 0.0009 |
| LC | Amblyodipsas microphthalma | 17 | 10.3418889 | 0.0009 |
| LC | Amblyodipsas polylepis | 127 | 10.8923983 | 0.0009 |
| LC | Aparallactus capensis | 409 | 10.8284276 | 0.0009 |
| LC | Aparallactus lunulatus | 101 | 11.608846 | 0.0009 |
| LC | Aspidelaps lubricus | 53 | 12.6621711 | 0.0009 |
| LC | Aspidelaps scutatus | 78 | 12.6546019 | 0.0009 |
| LC | Gerrhosaurus auritus | 5 | 22.6721289 | 0.0009 |
| LC | Atractaspis bibronii | 244 | 11.7364731 | 0.0009 |
| LC | Bitis arietans | 1083 | 18.6648655 | 0.0009 |
| LC | Bitis gabonica | 175 | 10.3076273 | 0.0009 |
| LC | Boaedon capensis | 536 | 8.86561803 | 0.0009 |
| LC | Boaedon fuliginosus | 467 | 8.08950819 | 0.0009 |
| LC | Broadleysaurus major | 113 | 50.2440279 | 0.0009 |

1. Appendix Sl

| | | | | |
|---|---|---|---|---|
| LC | Hemidactylus mercatorius | 75 | 13.9473209 | 0.0009 |
| LC | Causus defilippii | 153 | 10.3142583 | 0.0009 |
| LC | Causus rhombeatus | 389 | 10.5639694 | 0.0009 |
| LC | Chamaeleo dilepis | 769 | 14.3912223 | 0.0009 |
| LC | Homopholis arnoldi | 11 | 18.9483566 | 0.0009 |
| LC | Chirindia swynnertoni | 5 | 15.3767068 | 0.0009 |
| LC | Chondrodactylus bibronii | 321 | 25.174916 | 0.0009 |
| LC | Chondrodactylus turneri | 370 | 21.3148613 | 0.0009 |
| LC | Cordylus meculae | 2 | 14.0782866 | 0.0009 |
| LC | Cordylus tropidosternum | 73 | 14.1508862 | 0.0009 |
| LC | Cordylus vittifer | 316 | 15.7106586 | 0.0009 |
| LC | Crocodylus niloticus | 664 | 11.8369907 | 0.0009 |
| LC | Crotaphopeltis hotamboeia | 757 | 7.63872682 | 0.0009 |
| LC | Cryptoblepharus africanus | 31 | 6.52836438 | 0.0009 |
| LC | Cryptoblepharus boutonii | 52 | 6.80304182 | 0.0009 |
| LC | Cryptoblepharus caudatus | 1 | 7.89551391 | 0.0009 |
| LC | Lygodactylus capensis | 709 | 19.0190924 | 0.0009 |
| LC | Dalophia pistillum | 6 | 12.2120855 | 0.0009 |
| LC | Dasypeltis medici | 37 | 6.57294071 | 0.0009 |
| LC | Dasypeltis scabra | 727 | 6.90779236 | 0.0009 |
| LC | Dendroaspis angusticeps | 85 | 9.26844386 | 0.0009 |
| LC | Dermochelys coriacea | 2233 | 61.6865663 | 0.0009 |
| LC | Dipsadoboa aulica | 60 | 6.79166777 | 0.0009 |
| LC | Dispholidus typus | 656 | 10.1028213 | 0.0009 |
| LC | Duberria lutrix | 256 | 12.9189795 | 0.0009 |
| LC | Duberria variegata | 35 | 12.9023013 | 0.0009 |
| LC | Elapsoidea boulengeri | 26 | 8.53915597 | 0.0009 |
| LC | Elapsoidea nigra | 10 | 9.62241687 | 0.0009 |
| LC | Elapsoidea sundevallii | 107 | 7.72955569 | 0.0009 |
| LC | Elasmodactylus tetensis | 8 | 34.1293537 | 0.0009 |
| LC | Monopeltis decosteri | 1 | 10.8636452 | 0.0009 |
| LC | Gastropholis vittata | 4 | 17.9920746 | 0.0009 |
| LC | Geocalamus modestus | 2 | 15.0766068 | 0.0009 |
| LC | Rhamphiophis rostratus | 82 | 13.6894695 | 0.0009 |
| LC | Gerrhosaurus flavigularis | 527 | 19.6155436 | 0.0009 |
| LC | Gerrhosaurus nigrolineatus | 135 | 19.5025971 | 0.0009 |
| LC | Gonionotophis capensis | 110 | 11.0113275 | 0.0009 |
| LC | Gonionotophis nyassae | 70 | 12.4429374 | 0.0009 |
| LC | Heliobolus lugubris | 291 | 17.9179896 | 0.0009 |
| LC | Hemidactylus mabouia | 1558 | 13.9664873 | 0.0009 |
| LC | Nucras boulengeri | 41 | 27.8547723 | 0.0009 |
| LC | Hemidactylus platycephalus | 116 | 15.7208248 | 0.0009 |
| LC | Hemirhagerrhis nototaenia | 98 | 10.9428581 | 0.0009 |

1. Appendix Sl

| | | | | |
|---|---|---|---|---|
| LC | Holaspis laevis | 9 | 24.3606077 | 0.0009 |
| LC | Pachydactylus kobosensis | 8 | 15.8647604 | 0.0009 |
| LC | Homopholis walbergii | 225 | 19.3488888 | 0.0009 |
| LC | Hydrophis platurus | 429 | 5.05417272 | 0.0009 |
| LC | Ichnotropis capensis | 115 | 19.2853507 | 0.0009 |
| LC | Kinixys belliana | 133 | 16.9561143 | 0.0009 |
| LC | Lamprophis guttatus | 74 | 13.3502726 | 0.0009 |
| LC | Leptotyphlops incognitus | 86 | 13.5717622 | 0.0009 |
| LC | Leptotyphlops scutifrons | 303 | 14.0501217 | 0.0009 |
| LC | Lycodonomorphus leleupi | 2 | 10.7962499 | 0.0009 |
| LC | Lycophidion capense | 422 | 11.9071897 | 0.0009 |
| LC | Lycophidion semiannule | 1 | 10.539249 | 0.0009 |
| LC | Lycophidion variegatum | 19 | 10.2342496 | 0.0009 |
| LC | Platysaurus torquatus | 3 | 18.9825522 | 0.0009 |
| LC | Lygodactylus chobiensis | 37 | 22.1522193 | 0.0009 |
| LC | Lygodactylus conradti | 5 | 17.7925426 | 0.0009 |
| LC | Lygodactylus grotei | 29 | 18.8868432 | 0.0009 |
| LC | Lygodactylus picturatus | 101 | 19.8896873 | 0.0009 |
| LC | Prosymna stuhlmanni | 98 | 12.9923424 | 0.0009 |
| LC | Lygodactylus rex | 4 | 20.4779144 | 0.0009 |
| LC | Matobosaurus validus | 190 | 32.3180865 | 0.0009 |
| LC | Mecistops cataphractus | 35 | 18.9064621 | 0.0009 |
| LC | Meizodon semiornatus | 63 | 7.66537147 | 0.0009 |
| LC | Melanoseps ater | 12 | 10.5422361 | 0.0009 |
| LC | Meroles squamulosus | 235 | 26.7068925 | 0.0009 |
| LC | Mochlus afer | 94 | 7.56667431 | 0.0009 |
| LC | Mochlus sundevalli | 217 | 7.5744663 | 0.0009 |
| LC | Pseudocordylus melanotus | 120 | 14.0546886 | 0.0009 |
| LC | Monopeltis sphenorhynchus | 25 | 10.9809412 | 0.0009 |
| LC | Myriopholis ionidesi | 3 | 17.9855835 | 0.0009 |
| LC | Pelusios sinuatus | 163 | 43.9249291 | 0.0009 |
| LC | Naja annulifera | 180 | 6.53094604 | 0.0009 |
| LC | Naja melanoleuca | 265 | 9.27208738 | 0.0009 |
| LC | Naja mossambica | 304 | 6.16424155 | 0.0009 |
| LC | Naja naja | 125 | 6.13336373 | 0.0009 |
| LC | Natriciteres olivacea | 95 | 12.3894525 | 0.0009 |
| LC | Natriciteres sylvatica | 17 | 12.0482236 | 0.0009 |
| LC | Scelotes mirus | 81 | 14.0655079 | 0.0009 |
| LC | Nucras caesicaudata | 5 | 15.7742426 | 0.0009 |
| LC | Nucras holubi | 106 | 16.6946531 | 0.0009 |
| LC | Nucras ornata | 162 | 17.8386657 | 0.0009 |
| LC | Smaug warreni | 36 | 14.2695138 | 0.0009 |
| LC | Pachydactylus punctatus | 324 | 17.2094034 | 0.0009 |

1. Appendix Sl

| | | | | |
|---|---|---|---|---|
| LC | Proscelotes eggeli | 3 | 14.1005612 | 0.0009 |
| LC | Pelusios subniger | 35 | 43.9249291 | 0.0009 |
| LC | Philothamnus angolensis | 54 | 7.88709265 | 0.0009 |
| LC | Philothamnus hoplogaster | 246 | 8.22240408 | 0.0009 |
| LC | Philothamnus natalensis | 146 | 8.48472278 | 0.0009 |
| LC | Philothamnus punctatus | 46 | 7.52171122 | 0.0009 |
| LC | Philothamnus semivariegatus | 501 | 8.12080963 | 0.0009 |
| LC | Platysaurus intermedius | 174 | 19.1759097 | 0.0009 |
| LC | Platysaurus lebomboensis | 19 | 18.4996388 | 0.0009 |
| LC | Platysaurus maculatus | 8 | 19.0572753 | 0.0009 |
| LC | Trachylepis lacertiformis | 21 | 8.97082573 | 0.0009 |
| LC | Proatheris superciliaris | 6 | 16.1401697 | 0.0009 |
| LC | Trachylepis margaritifera | 186 | 11.522516 | 0.0009 |
| LC | Python sebae | 242 | 8.93192787 | 0.0009 |
| LC | Prosymna janii | 12 | 14.4111067 | 0.0009 |
| LC | Trachylepis striata | 818 | 8.54744314 | 0.0009 |
| LC | Psammophis angolensis | 73 | 10.1998486 | 0.0009 |
| LC | Psammophis mossambicus | 243 | 6.68769545 | 0.0009 |
| LC | Psammophis orientalis | 30 | 7.42834913 | 0.0009 |
| LC | Psammophis phillipsii | 218 | 8.61572817 | 0.0009 |
| LC | Psammophis sibilans | 122 | 6.74220577 | 0.0009 |
| LC | Psammophis subtaeniatus | 247 | 7.55282831 | 0.0009 |
| LC | Psammophylax tritaeniatus | 180 | 7.09522517 | 0.0009 |
| LC | Pseudaspis cana | 397 | 24.0672712 | 0.0009 |
| LC | Zygaspis violacea | 4 | 13.0629679 | 0.0009 |
| LC | Python natalensis | 290 | 7.63540677 | 0.0009 |
| LC | Stigmochelys pardalis | 719 | 26.6481397 | 0.0009 |
| LC | Tarentola mauritanica | 3364 | 14.3622783 | 0.0009 |
| LC | Telescopus semiannulatus | 246 | 7.99203361 | 0.0009 |
| LC | Tetradactylus ellenbergeri | 7 | 20.2408395 | 0.0009 |
| LC | Thelotornis capensis | 234 | 7.13106871 | 0.0009 |
| LC | Thelotornis mossambicanus | 15 | 7.23525654 | 0.0009 |
| LC | Thelotornis usambaricus | 9 | 7.25976167 | 0.0009 |
| LC | Trachylepis boulengeri | 11 | 10.2087735 | 0.0009 |
| LC | Trachylepis casuarinae | 1 | 10.2994584 | 0.0009 |
| LC | Scelotes mossambicus | 65 | 12.3535257 | 0.0009 |
| LC | Trachylepis homalocephala | 117 | 12.5565142 | 0.0009 |
| LC | Smaug mossambicus | 2 | 14.292828 | 0.0009 |
| LC | Trachylepis maculilabris | 282 | 9.94183744 | 0.0009 |
| LC | Trachylepis varia | 1013 | 12.1482814 | 0.0009 |
| LC | Trachylepis megalura | 40 | 9.74964775 | 0.0009 |
| LC | Trachylepis quinquetaeniata | 307 | 11.6431806 | 0.0009 |
| LC | Varanus exanthematicus | 194 | 10.9144628 | 0.0009 |

1. Appendix Sl

| | | | | |
|---|---|---|---|---|
| LC | Varanus niloticus | 919 | 12.8765504 | 0.0009 |
| LC | Trioceros melleri | 24 | 15.478645 | 0.0009 |
| LC | Varanus albigularis | 492 | 8.77015304 | 0.0009 |
| LC | Zygaspis vandami | 30 | 14.2465625 | 0.0009 |
| LC | Trachylepis depressa | 51 | 9.35699139 | 0.0009 |
| LC | Xenocalamus transvaalensis | 9 | 9.26191215 | 0.0009 |
| LC | Xenocalamus bicolor | 54 | 9.12475569 | 0.0009 |
| NT | Nadzikambia baylissi | 1 | 16.545787 | 0.0071 |
| NT | Rhampholeon maspictus | 1 | 18.3623304 | 0.0071 |
| NT | Lygodactylus regulus | 1 | 20.3898196 | 0.0071 |
| VU | Rhampholeon marshalli | 6 | 23.1364434 | 0.0513 |
| VU | Rhampholeon nebulauctor | 1 | 18.3946179 | 0.0513 |
| VU | Caretta caretta | 3584 | 34.1570163 | 0.0513 |

Table S2: Mammals species list, ED: Evolutionary distinctiveness, Range in number of 0.1-degree units, ER: Extinction risk.

| status | species | ED | Range | ER |
|---|---|---|---|---|
| DD | Mus neavei | 4.37900311 | 9 | 0.0513 |
| DD | Elephantulus fuscus | 13.944762 | 3 | 0.0513 |
| EN | Kerivoula africana | 4.23260653 | 2 | 0.4276 |
| EN | Lycaon pictus | 3.8718321 | 308 | 0.4276 |
| EN | Paraxerus vincenti | 7.29053014 | 3 | 0.4276 |
| LC | Acomys spinosissimus | 5.49607863 | 118 | 0.0009 |
| LC | Aepyceros melampus | 8.58036658 | 893 | 0.0009 |
| LC | Aethomys chrysophilus | 8.47579257 | 249 | 0.0009 |
| LC | Aethomys kaiseri | 8.17780417 | 125 | 0.0009 |
| LC | Alcelaphus buselaphus | 9.89034242 | 653 | 0.0009 |
| LC | Mellivora capensis | 17.8004848 | 223 | 0.0009 |
| LC | Atilax paludinosus | 13.5690442 | 232 | 0.0009 |
| LC | Beamys hindei | 17.2375484 | 35 | 0.0009 |
| LC | Calcochloris obtusirostris | 19.8361341 | 9 | 0.0009 |
| LC | Canis adustus | 3.45550887 | 280 | 0.0009 |
| LC | Canis lupus | 3.25661457 | 16308 | 0.0009 |
| LC | Genetta tigrina | 8.46129775 | 97 | 0.0009 |
| LC | Cephalophus natalensis | 5.95233965 | 87 | 0.0009 |
| LC | Cercopithecus mitis | 8.04067325 | 379 | 0.0009 |
| LC | Chaerephon ansorgei | 6.62934127 | 17 | 0.0009 |
| LC | Chaerephon pumilus | 6.02277361 | 338 | 0.0009 |
| LC | Chlorocebus aethiops | 4.97138325 | 368 | 0.0009 |
| LC | Chlorocebus pygerythrus | 4.99793255 | 597 | 0.0009 |
| LC | Civettictis civetta | 14.0716053 | 302 | 0.0009 |
| LC | Connochaetes taurinus | 8.55731471 | 685 | 0.0009 |

1. Appendix Sl

| | | | | |
|---|---|---|---|---|
| LC | Cricetomys gambianus | 13.4583229 | 366 | 0.0009 |
| LC | Crocidura cyanea | 5.10932601 | 45 | 0.0009 |
| LC | Crocidura fuscomurina | 4.87509502 | 91 | 0.0009 |
| LC | Crocidura hirta | 5.04849352 | 163 | 0.0009 |
| LC | Crocidura luna | 5.03971488 | 91 | 0.0009 |
| LC | Crocidura mariquensis | 5.04136371 | 33 | 0.0009 |
| LC | Crocidura nigrofusca | 5.32375744 | 37 | 0.0009 |
| LC | Crocidura olivieri | 5.08933906 | 434 | 0.0009 |
| LC | Crocidura silacea | 4.97918555 | 18 | 0.0009 |
| LC | Crocuta crocuta | 20.3873042 | 659 | 0.0009 |
| LC | Cryptomys hottentotus | 10.7807319 | 142 | 0.0009 |
| LC | Damaliscus lunatus | 9.49136675 | 310 | 0.0009 |
| LC | Dasymys incomtus | 7.66759514 | 243 | 0.0009 |
| LC | Dendrohyrax arboreus | 25.149054 | 123 | 0.0009 |
| LC | Dendromus melanotis | 13.3360865 | 102 | 0.0009 |
| LC | Dendromus messorius | 11.1102923 | 58 | 0.0009 |
| LC | Dendromus mystacalis | 10.74613 | 131 | 0.0009 |
| LC | Dendromus nyikae | 10.9032177 | 24 | 0.0009 |
| LC | Lemniscomys griselda | 7.51019245 | 124 | 0.0009 |
| LC | Elephantulus brachyrhynchus | 14.7194727 | 74 | 0.0009 |
| LC | Leptailurus serval | 9.8428388 | 213 | 0.0009 |
| LC | Epomophorus crypturus | 2.75060999 | 100 | 0.0009 |
| LC | Epomophorus gambianus | 2.75642778 | 209 | 0.0009 |
| LC | Epomophorus wahlbergi | 3.2984082 | 237 | 0.0009 |
| LC | Eptesicus hottentotus | 6.04351973 | 16 | 0.0009 |
| LC | Philantomba monticola | 9.00557633 | 209 | 0.0009 |
| LC | Felis silvestris | 7.26793166 | 5171 | 0.0009 |
| LC | Galago moholi | 14.5201612 | 111 | 0.0009 |
| LC | Genetta angolensis | 7.34216083 | 29 | 0.0009 |
| LC | Genetta genetta | 8.4780498 | 4611 | 0.0009 |
| LC | Genetta maculata | 6.99268524 | 336 | 0.0009 |
| LC | Mops condylurus | 7.48908461 | 188 | 0.0009 |
| LC | Gerbilliscus afra | 8.00286662 | 39 | 0.0009 |
| LC | Gerbilliscus inclusus | 7.79515324 | 6 | 0.0009 |
| LC | Gerbilliscus leucogaster | 7.99344087 | 337 | 0.0009 |
| LC | Gerbillurus paeba | 8.02841005 | 184 | 0.0009 |
| LC | Mus musculus | 3.89501752 | 18642 | 0.0009 |
| LC | Glauconycteris variegata | 5.49695142 | 45 | 0.0009 |
| LC | Grammomys cometes | 7.90622925 | 7 | 0.0009 |
| LC | Grammomys dolichurus | 8.14431173 | 314 | 0.0009 |
| LC | Grammomys macmillani | 7.70597813 | 50 | 0.0009 |
| LC | Graphiurus microtis | 13.9841324 | 47 | 0.0009 |
| LC | Graphiurus murinus | 14.2292683 | 239 | 0.0009 |

1. Appendix Sl

| | | | | |
|---|---|---|---|---|
| LC | Heliophobius argenteocinereus | 17.8538798 | 55 | 0.0009 |
| LC | Heliosciurus mutabilis | 7.75988499 | 46 | 0.0009 |
| LC | Herpestes ichneumon | 10.706005 | 1009 | 0.0009 |
| LC | Heterohyrax brucei | 26.1482966 | 161 | 0.0009 |
| LC | Oreotragus oreotragus | 8.53759707 | 342 | 0.0009 |
| LC | Hipposideros caffer | 9.68301425 | 368 | 0.0009 |
| LC | Hipposideros ruber | 10.1253708 | 204 | 0.0009 |
| LC | Otomys angoniensis | 7.37779984 | 82 | 0.0009 |
| LC | Hippotragus equinus | 8.07972884 | 717 | 0.0009 |
| LC | Hippotragus niger | 8.07666356 | 254 | 0.0009 |
| LC | Hystrix africaeaustralis | 9.10941315 | 248 | 0.0009 |
| LC | Ictonyx striatus | 9.68272387 | 157 | 0.0009 |
| LC | Papio ursinus | 5.85614827 | 506 | 0.0009 |
| LC | Kobus ellipsiprymnus | 12.01706 | 908 | 0.0009 |
| LC | Laephotis botswanae | 7.48001464 | 15 | 0.0009 |
| LC | Paraxerus palliatus | 7.2125445 | 43 | 0.0009 |
| LC | Lemniscomys rosalia | 7.43441124 | 111 | 0.0009 |
| LC | Pedetes capensis | 35.301966 | 167 | 0.0009 |
| LC | Pelomys fallax | 9.95375558 | 201 | 0.0009 |
| LC | Petrodromus tetradactylus | 21.7638216 | 77 | 0.0009 |
| LC | Mastomys coucha | 8.06242418 | 88 | 0.0009 |
| LC | Mastomys natalensis | 8.21453699 | 1112 | 0.0009 |
| LC | Tadarida aegyptiaca | 10.5910484 | 91 | 0.0009 |
| LC | Miniopterus africanus | 8.66371063 | 26 | 0.0009 |
| LC | Miniopterus fraterculus | 9.4426477 | 36 | 0.0009 |
| LC | Miniopterus inflatus | 8.6935478 | 64 | 0.0009 |
| LC | Miniopterus natalensis | 8.45398708 | 103 | 0.0009 |
| LC | Potamochoerus porcus | 13.9767897 | 151 | 0.0009 |
| LC | Praomys delectorum | 7.93478413 | 55 | 0.0009 |
| LC | Mops niveiventer | 7.52065938 | 13 | 0.0009 |
| LC | Mungos mungo | 10.0504503 | 259 | 0.0009 |
| LC | Mus minutoides | 4.34890824 | 273 | 0.0009 |
| LC | Mus musculoides | 4.43255767 | 543 | 0.0009 |
| LC | Rattus norvegicus | 4.98697001 | 13094 | 0.0009 |
| LC | Rattus rattus | 4.49261596 | 7696 | 0.0009 |
| LC | Mus triton | 4.35216255 | 196 | 0.0009 |
| LC | Myotis bocagii | 5.25976838 | 93 | 0.0009 |
| LC | Myotis tricolor | 4.52970813 | 42 | 0.0009 |
| LC | Neoromicia capensis | 7.28812566 | 203 | 0.0009 |
| LC | Neoromicia nana | 7.95107559 | 370 | 0.0009 |
| LC | Nycteris grandis | 12.399647 | 60 | 0.0009 |
| LC | Nycteris macrotis | 10.4608168 | 144 | 0.0009 |
| LC | Nycteris thebaica | 11.1153216 | 362 | 0.0009 |

1. Appendix Sl

| | | | | |
|---|---|---|---|---|
| LC | Nycteris woodi | 11.3456956 | 13 | 0.0009 |
| LC | Rhinolophus hildebrandtii | 2.94154624 | 115 | 0.0009 |
| LC | Otolemur crassicaudatus | 12.9397862 | 161 | 0.0009 |
| LC | Otolemur garnettii | 12.9612807 | 79 | 0.0009 |
| LC | Rhinolophus swinnyi | 3.01252648 | 28 | 0.0009 |
| LC | Ourebia ourebi | 8.47232876 | 387 | 0.0009 |
| LC | Rhynchogale melleri | 13.7356565 | 5 | 0.0009 |
| LC | Rousettus aegyptiacus | 2.72563273 | 304 | 0.0009 |
| LC | Papio cynocephalus | 5.75762254 | 202 | 0.0009 |
| LC | Sauromys petrophilus | 12.5456861 | 14 | 0.0009 |
| LC | Paraxerus cepapi | 7.21028962 | 316 | 0.0009 |
| LC | Paraxerus flavovittis | 6.99797652 | 9 | 0.0009 |
| LC | Scotophilus nigrita | 5.44685452 | 60 | 0.0009 |
| LC | Scotophilus viridis | 5.15073616 | 76 | 0.0009 |
| LC | Steatomys krebsii | 11.3478296 | 22 | 0.0009 |
| LC | Steatomys pratensis | 11.0667928 | 145 | 0.0009 |
| LC | Suncus megalura | 7.6702364 | 89 | 0.0009 |
| LC | Phacochoerus aethiopicus | 17.3689864 | 170 | 0.0009 |
| LC | Phacochoerus africanus | 17.2210691 | 1021 | 0.0009 |
| LC | Redunca arundinum | 9.61598224 | 369 | 0.0009 |
| LC | Pipistrellus hesperidus | 7.96704072 | 63 | 0.0009 |
| LC | Pipistrellus rueppellii | 8.74158316 | 74 | 0.0009 |
| LC | Poecilogale albinucha | 12.5685394 | 49 | 0.0009 |
| LC | Potamochoerus larvatus | 14.0157959 | 226 | 0.0009 |
| LC | Thryonomys gregorianus | 13.9785765 | 43 | 0.0009 |
| LC | Thryonomys swinderianus | 13.9785765 | 252 | 0.0009 |
| LC | Procavia capensis | 25.7683964 | 611 | 0.0009 |
| LC | Pronolagus rupestris | 14.6013178 | 15 | 0.0009 |
| LC | Raphicerus campestris | 7.03152568 | 579 | 0.0009 |
| LC | Raphicerus sharpei | 7.03237554 | 49 | 0.0009 |
| LC | Uranomys ruddi | 8.65821115 | 120 | 0.0009 |
| LC | Sylvicapra grimmia | 7.10003979 | 777 | 0.0009 |
| LC | Rhynchocyon cirnei | 24.6362768 | 43 | 0.0009 |
| LC | Rhabdomys pumilio | 10.6594785 | 420 | 0.0009 |
| LC | Rhinolophus blasii | 2.90303409 | 102 | 0.0009 |
| LC | Rhinolophus capensis | 3.0327053 | 16 | 0.0009 |
| LC | Rhinolophus clivosus | 2.80837458 | 171 | 0.0009 |
| LC | Rhinolophus darlingi | 3.03161386 | 36 | 0.0009 |
| LC | Scotophilus dinganii | 5.13529237 | 221 | 0.0009 |
| LC | Rhinolophus denti | 3.00704787 | 21 | 0.0009 |
| LC | Rhinolophus fumigatus | 2.94604361 | 130 | 0.0009 |
| LC | Tragelaphus scriptus | 9.72977769 | 883 | 0.0009 |
| LC | Rhinolophus landeri | 2.82378979 | 127 | 0.0009 |

1. Appendix Sl

| | | | | |
|---|---|---|---|---|
| LC | Rhinolophus simulator | 2.94188611 | 69 | 0.0009 |
| LC | Saccostomus campestris | 15.887553 | 311 | 0.0009 |
| LC | Syncerus caffer | 10.6257933 | 1466 | 0.0009 |
| LC | Scotoecus hirundo | 9.70037818 | 70 | 0.0009 |
| LC | Tadarida fulminans | 8.88613185 | 14 | 0.0009 |
| LC | Triaenops persicus | 13.7136535 | 42 | 0.0009 |
| LC | Taphozous mauritianus | 16.8536457 | 192 | 0.0009 |
| LC | Tragelaphus strepsiceros | 9.20402836 | 824 | 0.0009 |
| LC | Tragelaphus angasii | 9.39239649 | 177 | 0.0009 |
| LC | Thallomys paedulcus | 8.41305236 | 55 | 0.0009 |
| LC | Tadarida lobata | 9.02176784 | 9 | 0.0009 |
| NT | Equus quagga | 12.0503348 | 1395 | 0.0071 |
| NT | Hipposideros vittatus | 10.6466515 | 32 | 0.0071 |
| NT | Aonyx capensis | 5.46639153 | 154 | 0.0071 |
| NT | Eidolon helvum | 6.6856344 | 457 | 0.0071 |
| NT | Rhinolophus deckenii | 3.0458783 | 24 | 0.0071 |
| NT | Miniopterus schreibersii | 8.25713051 | 3550 | 0.0071 |
| VU | Hippopotamus amphibius | 32.8903701 | 1239 | 0.0513 |
| VU | Giraffa camelopardalis | 16.4828711 | 651 | 0.0513 |
| VU | Panthera leo | 8.25922665 | 1359 | 0.0513 |
| VU | Panthera pardus | 8.26451043 | 980 | 0.0513 |
| VU | Carpitalpa arendsi | 17.8620023 | 1 | 0.0513 |
| VU | Loxodonta africana | 37.3127961 | 1612 | 0.0513 |

Table S3: Perfect score by number of areas triggering KBA status

| Number of KBAs | Perfect ranking |
|---|---|
| 1 | 1 |
| 1+2 | 3 |
| 1+2+3 | 6 |
| 1+2+3+4 | 10 |
| 1+2+3+4+5 | 15 |
| 1+2+3+4+5+6 | 21 |
| 1+2+3+4+5+6+7 | 28 |
| 1+2+3+4+5+6+7+8 | 36 |

1. Appendix Sl

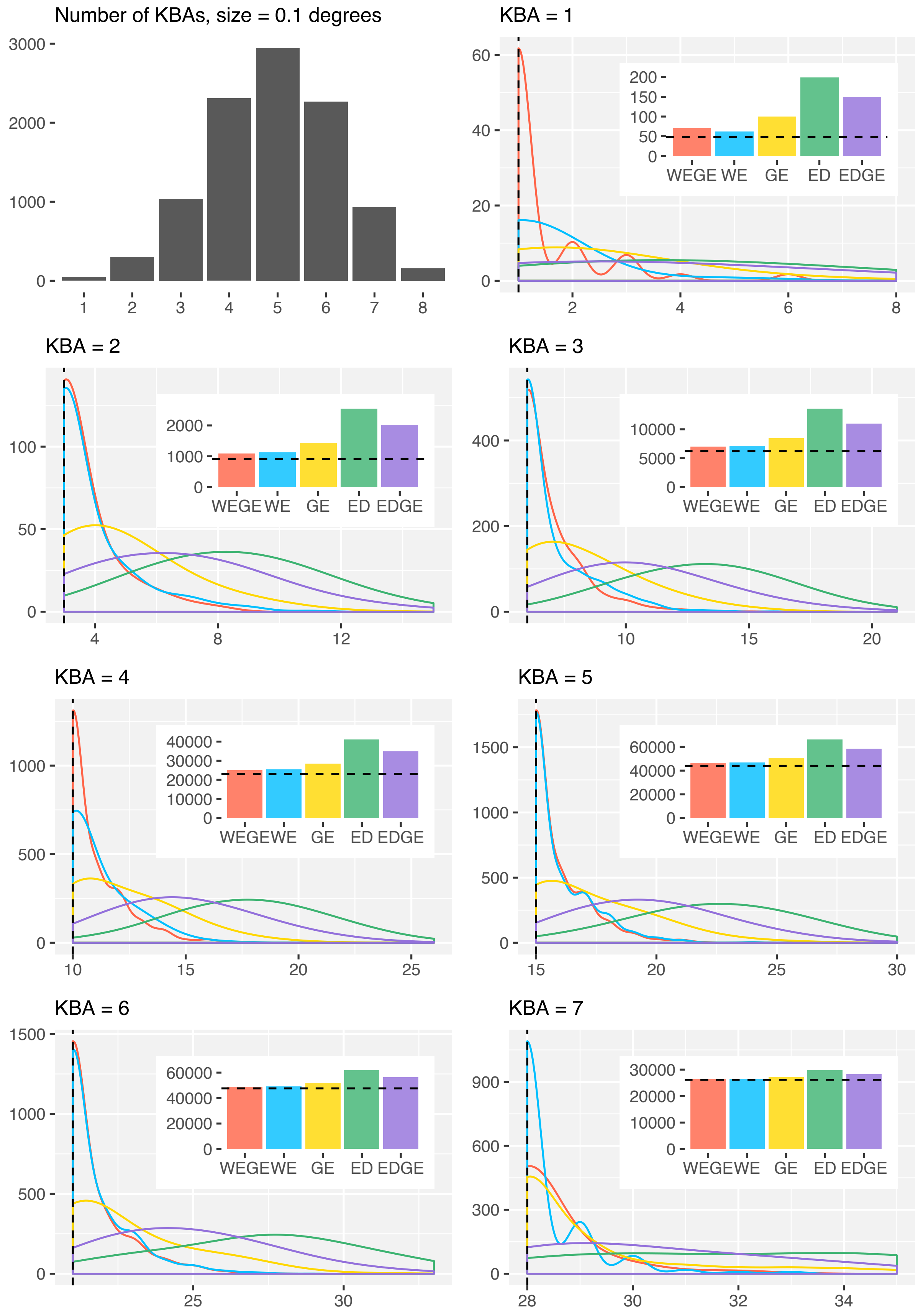


Figure S1: Reptile analysis for areas with 0.1 by 0.1 degrees of area. Density plot corresponds to the count of the different scores obtained inn each metric and bar chart corresponds to the sum of all scores in each index. Dashed line in both plots corresponds to the perfect score.

1. Appendix Sl

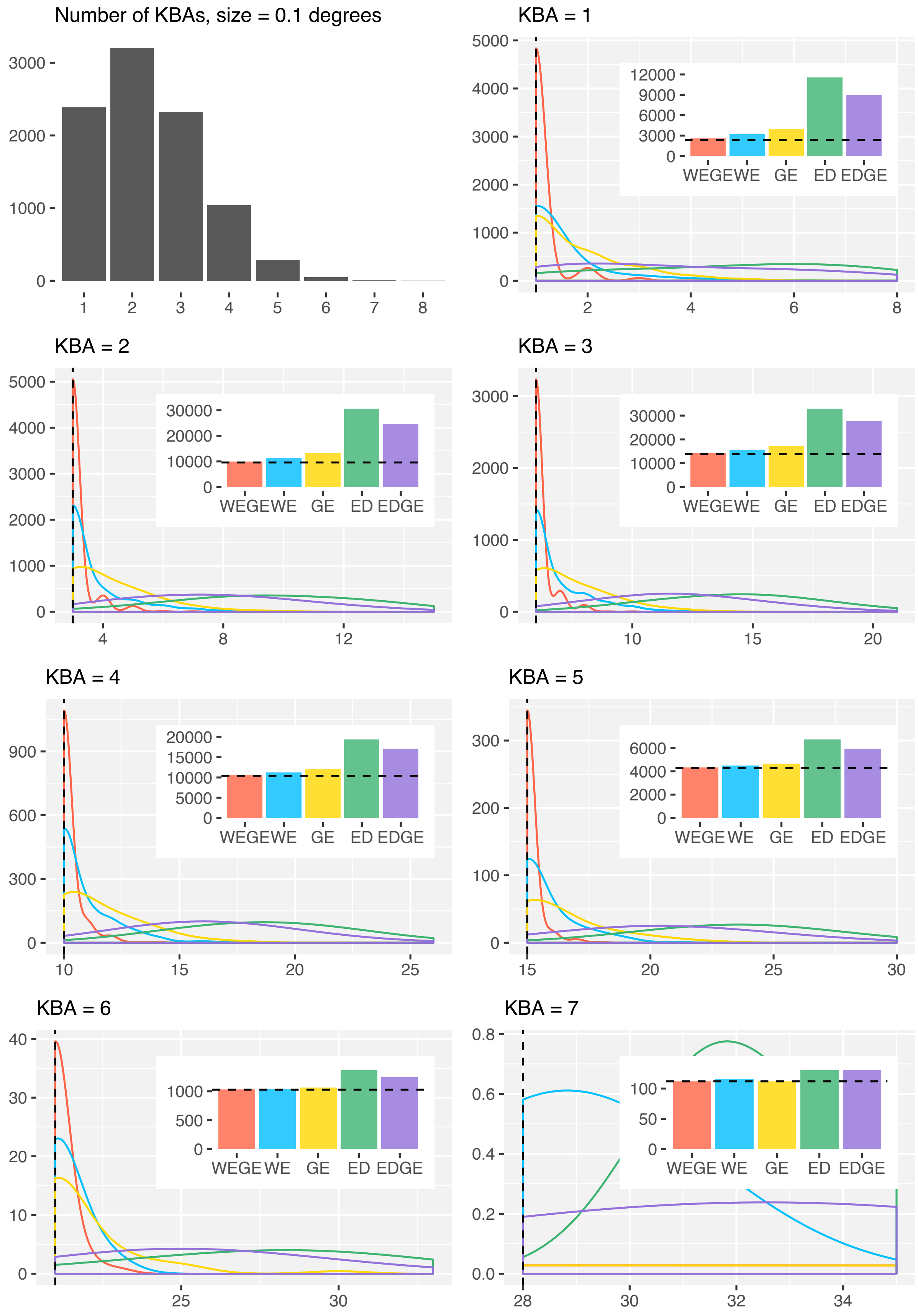


Figure 2: Mammal analysis for areas with 0.1 by 0.1 degrees of area. Density plot corresponds to the count of the different scores obtained inn each metric and bar chart corresponds to the sum of all scores in each index. Dashed line in both plots corresponds to the perfect score.

1. Appendix Sl

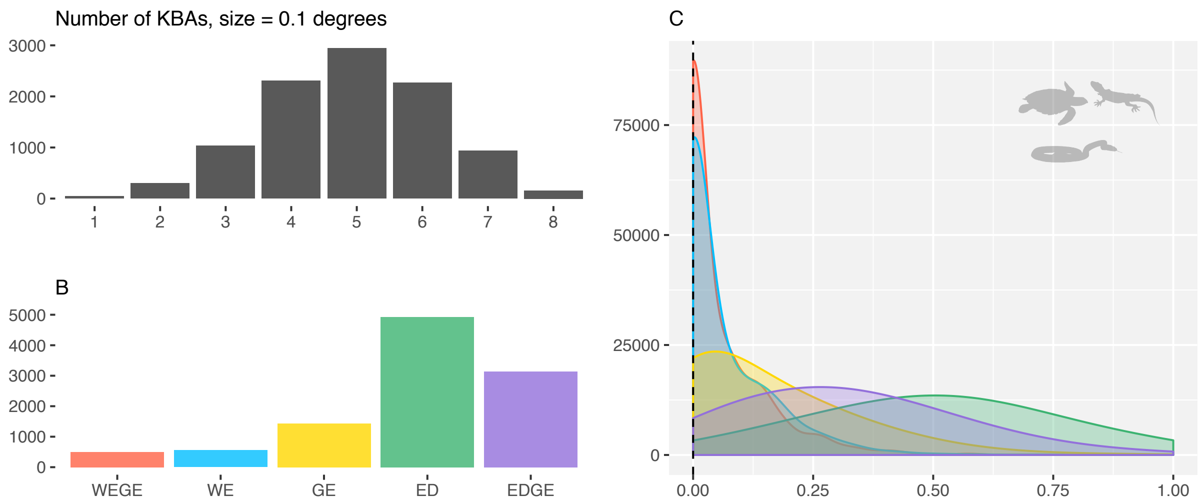


Figure S3: A. Number of areas triggering KBA status obtained by simulating 10 000 scenarios in reptile species' composition. B. Indices combined sum for all scenarios. C. Frequency of scores normalized between different number of KBAs. The figure shows that WEGE outperforms the other indices by both getting a smaller overall sum (B) and by having a higher density of values closer to 0 (C).

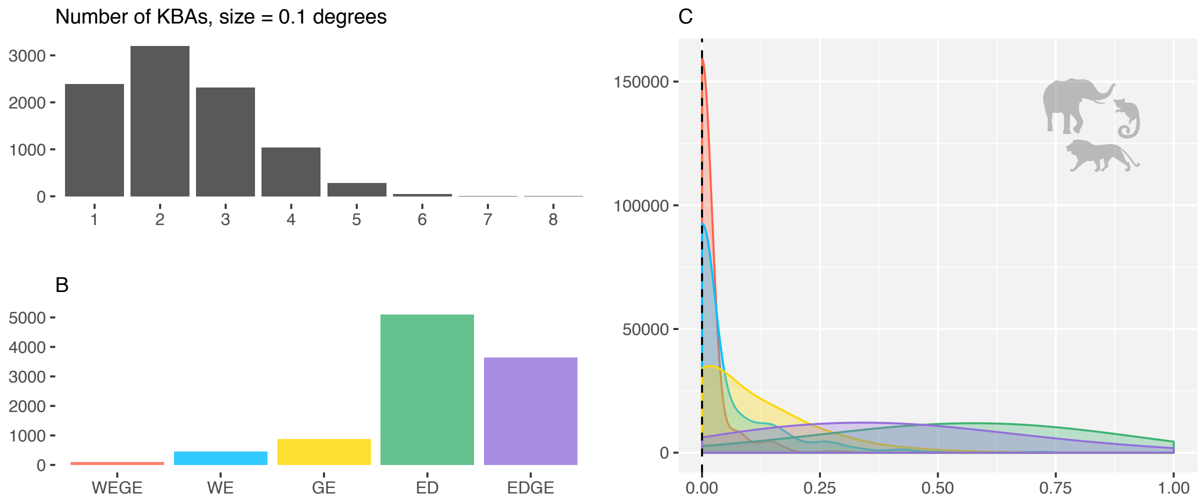


Figure S4: A. Number of areas triggering KBA status obtained by simulating 10 000 scenarios in mammal species' composition. B. Indices combined sum for all scenarios. C. Frequency of scores normalized between different number of KBAs. The figure shows that WEGE outperforms the other indices by both getting a smaller overall sum (B) and by having a higher density of values closer to 0 (C).

1. Appendix Sl

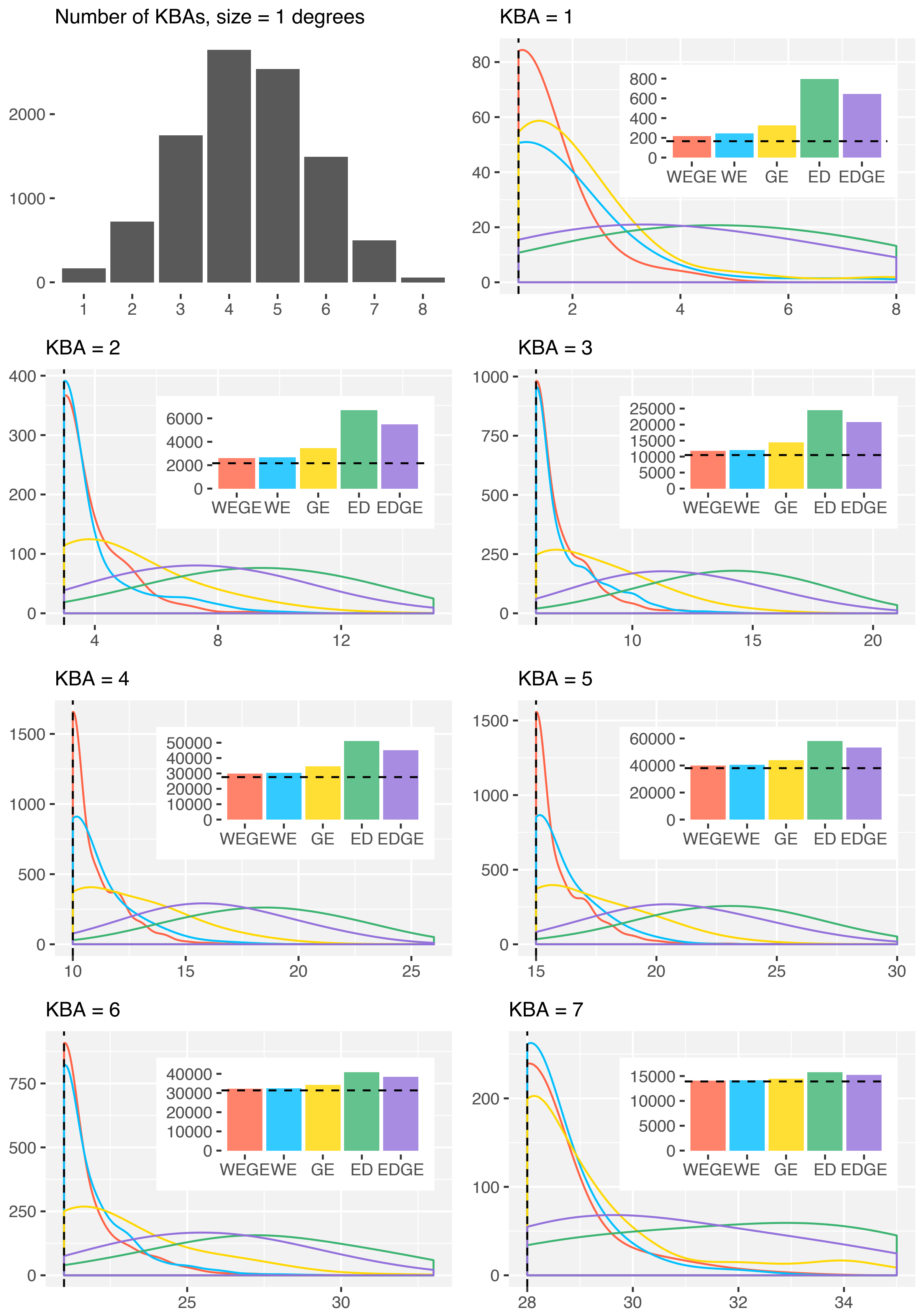


Figure S5: Reptile analysis for areas with 1 by 1 degrees of area. Density plot corresponds to the count of the different scores obtained inn each metric and bar chart corresponds to the sum of all scores in each index. Dashed line in both plots corresponds to the perfect score.

1. Appendix Sl

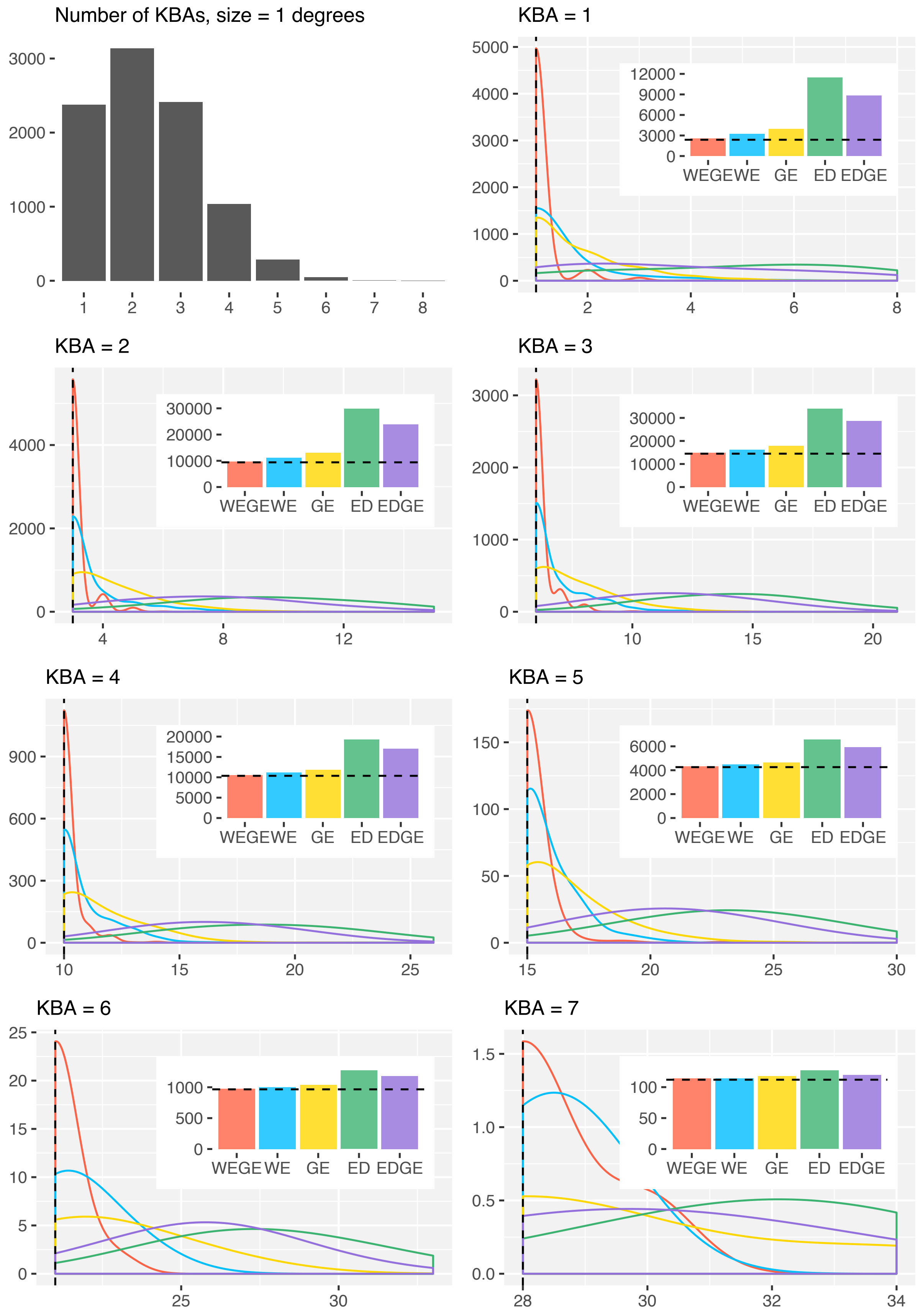


Figure S6: Mammal analysis for areas with 1 by 1 degrees of area. Density plot corresponds to the count of the different scores obtained inn each metric and bar chart corresponds to the sum of all scores in each index. Dashed line in both plots corresponds to the perfect score.

1. Appendix Sl

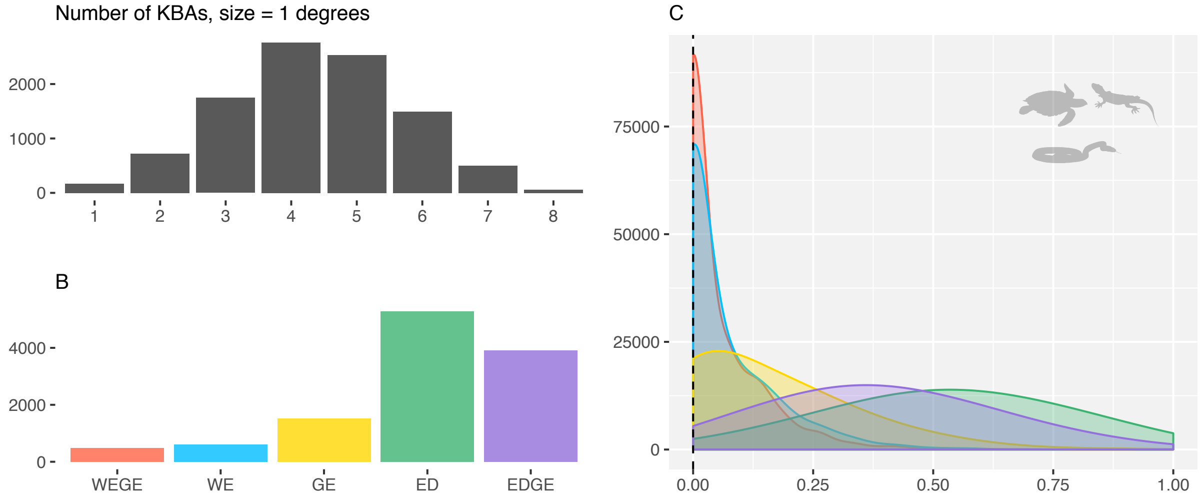


Figure S7: A. Number of areas triggering KBA status obtained by simulating 10 000 scenarios in reptile species' composition. B. Indices combined sum for all scenarios. C. Frequency of scores normalized between different number of KBAs. The figure shows that WEGE outperforms the other indices by both getting a smaller overall sum (B) and by having a higher density of values closer to 0 (C).

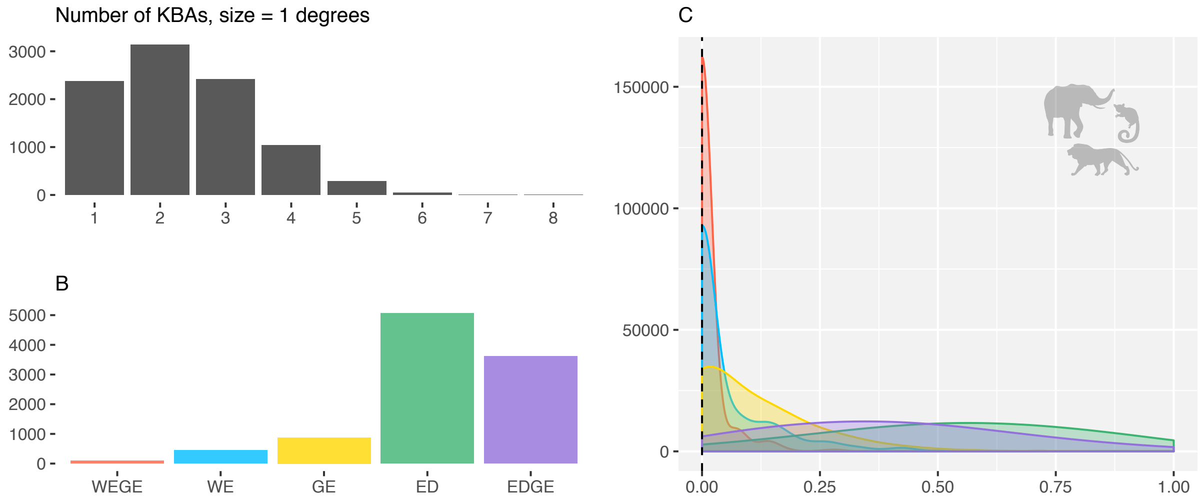


Figure S8: A. Number of areas triggering KBA status obtained by simulating 10 000 scenarios in mamal species' composition. B. Indices combined sum for all scenarios. C. Frequency of scores normalized between different number of KBAs. The figure shows that WEGE outperforms the other indices by both getting a smaller overall sum (B) and by having a higher density of values closer to 0 (C).
